# Regulation of the Balance between Concentric and Eccentric Cardiac Hypertrophy by a CDC14A-KMT5A Signaling Pathway

**DOI:** 10.64898/2026.02.16.706249

**Authors:** Xueyi Li, Jinliang Li, Yuliang Tan, Anne-Maj Samuelsson, Vi B. Nguyen, Ramesh V. Nair, Anne-Sophie Colombe, Dirk Grimm, Michael G. Rosenfeld, Michael S. Kapiloff

**Affiliations:** Stanford Cardiovascular Institute, Stanford University, Palo Alto, CA 94304; Department of Ophthalmology, Stanford University, Palo Alto, CA 94304; Department of Medicine, University of California, San Diego, 9500 Gilman Drive, La Jolla, CA 92093; Stanford Center for Genomics and Personalized Medicine, Stanford University School of Medicine, Stanford, CA, 94305; Department of Infectious Diseases/Virology, Section Viral Vector Technologies, Medical Faculty, BioQuant, Heidelberg University, Heidelberg, Germany; Department of Medicine (Cardiovascular Medicine), Stanford University, Palo Alto, CA 94304

**Keywords:** cardiac hypertrophy, signal transduction, CDC14A, KMT5A, H4K20me1, Dilated Cardiomyopathy

## Abstract

**Background:** Depending upon the type of pathological stress, the heart undergoes concentric or eccentric remodeling. This structural change is associated with diastolic and/or systolic ventricular dysfunction reflecting differentially altered cardiomyocyte morphology, ultrastructure, metabolism, contractility, and survival, as well as interstitial myocardial fibrosis. Despite an association of both concentric and eccentric remodeling with heart failure and sudden death, the molecular mechanisms resulting in abnormal cardiac geometry remain poorly understood. A better understanding of the basic mechanisms conferring these contrasting forms of remodeling should inform novel approaches to preserve normal cardiac structure and function in cardiovascular disease. The protein phosphatase Cell Division Cycle 14A (CDC14A) and its substrate the lysine methyltransferase KMT5A are identified herein as key regulators of the balance between concentric and eccentric pathological cardiac remodeling.

**Methods:** The regulation of adult rat ventricular myocyte morphology by CDC14A and KMT5A was studied *in vitro* following gain and loss of function by expression of wild-type and mutant proteins and RNA interference (RNAi). Epigenomic regulation by KMT5A was studied by mapping histone 4 lysine 20 mono-methylation (H4K20me1) modified chromatin sites and correlating them with gene transcription. Regulation of pathological cardiac remodeling *in vivo* was demonstrated by CDC14A and KMT5A RNAi using adeno-associated virus (AAV) mediated cardiomyocyte-specific small hairpin RNA (shRNA) expression in mice.

**Results:** CDC14A inhibited the growth in width of cultured adult myocytes stimulated by α-adrenergic receptor activation or by serum response factor. KMT5A was downregulated by CDC14A in cardiomyocytes and was required for myocyte growth in width. α-adrenergic stimulation of KMT5A-dependent H4K20 mono-methylation across transcription units correlated with regulation of gene transcription. Accordingly, AAV-expressed KMT5A shRNA induced eccentric remodeling and cardiac dysfunction in wild-type mice. Conversely, expression of Cdc14A shRNA improved systolic function and cardiac structure and inhibited pathological gene expression in the *Tpm1* E54K mouse with Dilated Cardiomyopathy.

**Conclusions:** CDC14A-KMT5A-dependent epigenomic regulation of gene transcription constitutes a molecular switch that determines concentric versus eccentric cardiac remodeling. These findings identify CDC14A as a potential therapeutic target for the treatment of dilated cardiomyopathy and other forms of heart failure with reduced ejection fraction.

**Clinical Perspective:** *What is new:*

- A function is identified for the first time for the protein phosphatase CDC14A in the heart, regulation of cardiomyocyte morphology and overall cardiac geometry in pathological cardiac remodeling.
- The lysine methyltransferase KMT5A is shown to mediate the effects of CDC14A in the adult cardiomyocyte by regulating H4K20 mono-methylation, such that reduced KMT5A expression promotes a phenotype resembling Dilated Cardiomyopathy.
- H4K20me1 epigenomic modification is identified as a regulator of cardiac structure and function.

*Clinical implications:*

- CDC14A loss of function experimentation *in vivo*, resulting in improved cardiac structure and function in a mouse model of Dilated Cardiomyopathy, suggests that CDC14A is a novel therapeutic target for heart failure with reduced ejection fraction.

## Introduction

The morphology and size of the heart are tightly regulated, such that deviations from the normal range in left ventricular (LV) shape and relative wall thickness (wall thickness to interior diameter ratio) are indicative of disease.^1^ Notably, altered cardiac geometry is the most outward manifestation of pathological cardiac remodeling, which includes altered myocyte contractility, metabolism, and survival, myocyte hypertrophy, myocardial inflammation, and interstitial fibrosis. However, structural cardiac remodeling not only is a marker of disease, but also results in abnormal ventricular mechanics, contributing to aberrant neurohormonal, paracrine, and autocrine cardiomyocyte stimulation and activation of mechanosensitive pathways.^1–3^ This activation of pathological intracellular signaling worsens remodeling, exacerbating diastolic and/or systolic cardiac dysfunction. Based upon the change in relative wall thickness, remodeling can be classified as concentric and eccentric, representing a spectrum of remodeling with differences in underlying cellular defects.^4^ Derivation of strategies to normalize relative wall thickness, improving both ventricular mechanics and associated cellular stress signaling, may provide new therapies to prevent or treat heart failure, a problem of major public health significance.

Although diverse signaling pathways have been shown to promote pathological cardiac remodeling, only extracellular signal-regulated kinase (ERK) signaling pathways have been demonstrated to influence the balance between concentric and eccentric cardiac hypertrophy.^4–6^ Activation of the ERK1/2 pathway by expression of a constitutively active mitogen-activated protein kinase kinase 1 (MEK1) or ERK1 transgene resulted in concentric cardiac hypertrophy, with cardiomyocytes characterized by an increased width:length (W:L) ratio, while, conversely, ERK1/2 double knock-out resulted in dilated cardiomyopathy (DCM) and elongated myocytes.^7,8^ We showed that the ERK effector p90 ribosomal S6 kinase type 3 (RSK3) is required for concentric cardiac hypertrophy, both in response to pressure overload and in a mouse model of Noonan’s Syndrome, a human cardiomyopathy resulting from activating mutations of the ERK1/2 pathway.^9,10^ Subsequently, we demonstrated that RSK3-dependent serum response factor (SRF) serine-103 (S103) phosphorylation promotes cardiomyocyte growth in width.^11^ Expression in mice of a SRF S103D phosphomimetic mutant induced mild concentric hypertrophy with interstitial fibrosis. Conversely, inhibition of RSK3 signaling by adeno-associated virus (AAV) -mediated expression of an A-kinase anchoring protein 6 (AKAP6/mAKAP)-derived RSK3 anchoring disruptor peptide prevented concentric hypertrophy and heart failure in response to short term and long term pressure overload, respectively.^11^ These results suggested that whereas therapeutic targeting of ERK1/2 signaling is complicated by the role of ERK1/2 in cardiomyocyte survival,^5^ inhibition of the downstream RSK3-SRF pathway might be useful in cardiac diseases featuring concentric pathological remodeling.

To determine the transcriptomic effects of activated ERK-RSK3-SRF signaling, we performed precision nuclear run-on sequencing (PRO-seq) to detect nascent pre-mRNA transcripts in adult rat ventricular myocytes.^11^ As described below, C*dc14a* transcription was inhibited by the ERK-RSK3-SRF pathway. CDC14A is a dual specificity tyrosine and serine/threonine protein phosphatase (Figure 1A),^12^ whose function in the heart has not heretofore been reported. We describe how CDC14A acts as a brake on cardiomyocyte growth in width, thereby determining eccentric versus concentric cardiac hypertrophy. In addition, the lysine methyltransferase KMT5A (also known as PR-SET7, Setd8, and Set8) is shown to mediate the effects of CDC14A in cardiomyocytes by epigenetically regulating H4K20 mono-methylation.

**Figure 1.**
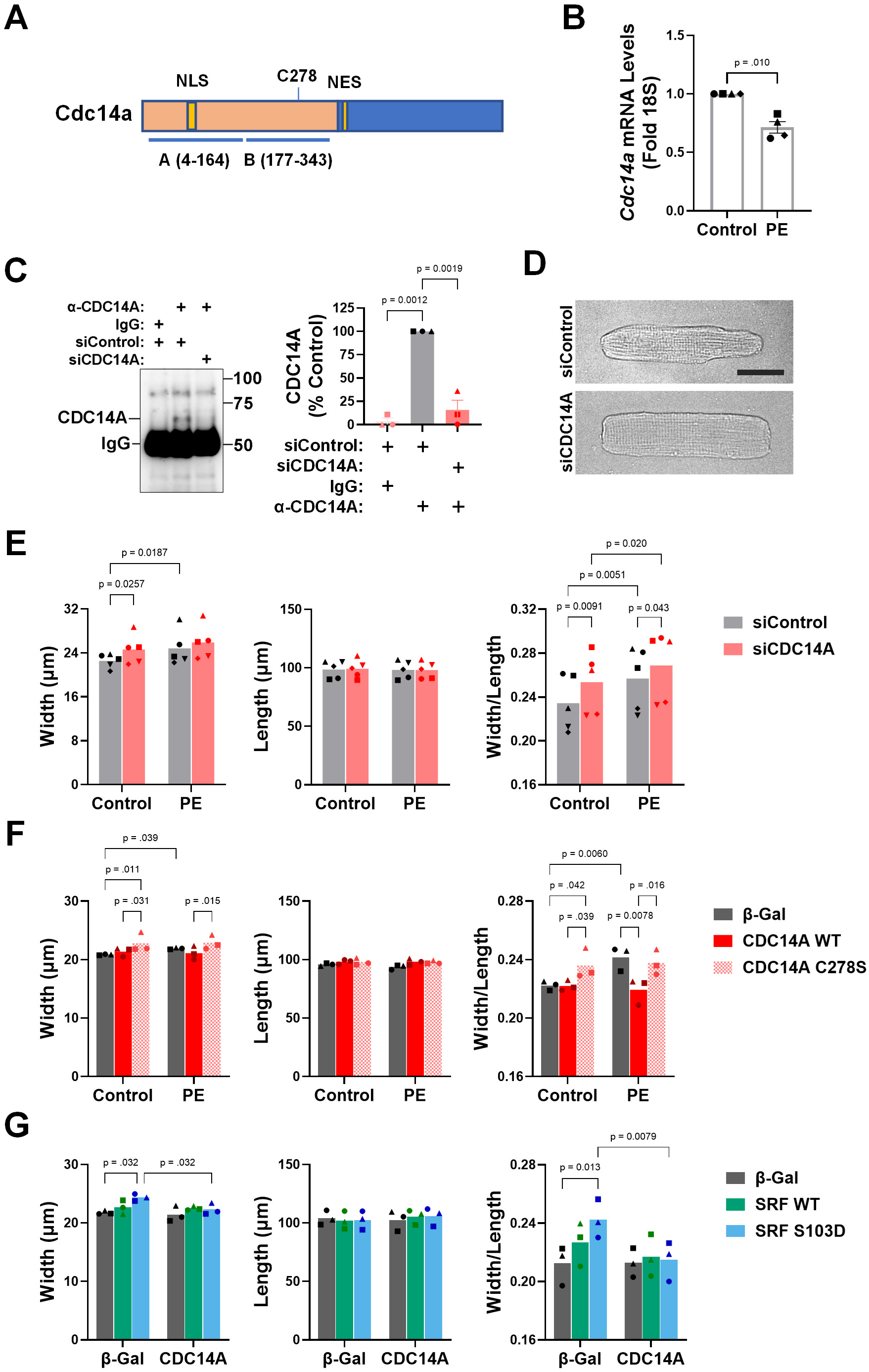
CDC14A regulates the growth in width of cultured adult rat ventricular myocytes. **A.** CDC14A contains a N-terminal phosphatase catalytic core (beige) which is divided into A and B domains conferring substrate specificity and catalytic activity, respectively.^12^ It has nuclear localization (NLS) and export (NES) signals.^16^ **B.** Adult rat ventricular myocytes were treated for 24 hours ± 20 µmol/L phenylephrine (PE) before total RNA isolation and qRT-PCR for *Cdc14a* and *18S* RNA levels. n = 4. Paired t-test. **C.** Neonatal rat ventricular myocytes transfected with control or CDC14a siRNA for 3 days before protein immunoprecipitation with preimmune or CDC14A goat antiserum and Western blot using an HRP-conjugated mouse anti-CDC14A antibody. Samples compared by 1-way ANOVA (matched for myocyte preparation) and Tukey’s post-hoc test; *n* = 3 separate myocyte preparations. **D,E.** Adult myocytes transfected for 24 hours with either CDC14A or control siRNA were then treated for 24 hours ± 20 µmol/L PE before measurement of myocyte width and length. *n* = 5 separate myocyte preparations. D – images of myocytes in minimal medium; bar – 25 µm. **F.** Adenoviral-transduced adult myocytes expressing Flag-tagged CDC14A wild-type (WT) or C278S catalytically inactive mutant or β-galactosidase (β-gal) control were treated for 24 hours ± 20 µmol/L PE, before measurement of individual myocyte width and length. *n* = 3 separate myocyte preparations. **G.** Adenoviral-transduced adult myocytes expressing SRF WT or S103D mutant and/or Flag-tagged CDC14A or β-galactosidase (β-gal) control were cultured in minimal media for 48 hours before measurement of width and length. *n* = 3 separate myocyte preparations. For E-G, ∼100 myocytes were measured per biological replicate sample. See Figure S1 for myocyte images. The mean values for each individual myocyte preparation are indicated by differently shaped symbols. *p*-values for comparison of mean width, length and width/length ratio are based on 2-way ANOVA with samples matched by myocyte preparation, with Fisher’s LSD (E) or Tukey’s Post-hoc testing (F,G).

## Methods

Detailed methods are provided in the Online Supplemental Material. All original data in this publication are available upon reasonable request to the corresponding authors. CUT&RUN and RNA-seq datasets are available at NCBI GEO GSE318470 and GSE318858, respectively.

### Animal Studies

All animal research was approved by the Institutional Animal Care and Use Committee (IACUC) at Stanford University. 2-month-old male Sprague–Dawley rats were used for isolation of adult rat ventricular myocytes. KMT5A loss of function was studied by intravenous administration of 10^11^ viral genomes (vg) AAV with the AAVMYO capsid^13^ into 6-week-old male wild-type C57BL/6NJ mice (Jackson Laboratory strain #005304). CDC14A loss of function was studied by intraperitoneal (IP) injection of 10^11^ vg serotype 9 AAV into 2-3-day-old male and female FVB/N-Tg(Myh6-Tpm1*E54K)67Dfw/MskfJ “TM54” mice (Jackson Laboratory strain #035610) for the study of DCM.^14^

### Statistical analysis

Statistics were computed using Prism 10 (Graphpad, San Diego, California). All data are expressed as mean ± s.e.m. The statistical test used for each experiment is indicated in the figure or table legend. Complete statistical methods are included in the Supplemental Material.

## Results

### Inhibited CDC14A Expression Promotes the Growth in Width of Adult Cardiomyocytes

In response to α-adrenergic receptor (αAR) stimulation, the ERK-RSK3-SRF pathway increased myocyte W:L ratio *in vitro*.^8,11^ We previously performed PRO-seq to detect nascent pre-mRNA transcripts in adult rat ventricular myocytes stimulated with the αAR agonist phenylephrine (PE) or expressing the hypertrophic phosphomimetic SRF S103D mutant protein (NCBI GEO accession number GSE134801).^11^ In that study, there was a significant correlation in the changes in gene transcription induced by PE stimulation for 1 hour and SRF S103D expression for 1 day, including 257 genes significantly altered in expression by both PE and SRF S103D (adjusted *p*-value < 0.05). Although the DNA binding of Ser-103 phosphorylated SRF is associated with gene activation,^11^ such that genes repressed by SRF S103D would be presumably indirectly regulated, we considered that an RNA interference screen would be a straightforward approach to identifying new relevant effector genes downstream of PE and SRF S103D that regulate myocyte morphology. Of the 96 genes repressed by both PE and SRF S103D expression in the PRO-seq study, one gene whose function had not previously been described in the heart was *Cdc14a*. PRO-seq detected two *Cdc14a* transcripts that encode CDC14A proteins with different C-terminal peptides. The transcription of NM_001134856 encoding the 67 kDa variant 1 was repressed by PE with a log_2_FC = -0.35 (adjusted *p*-value = 2.6 × 10^−4^) and by SRF S103D expression with a log_2_FC = -0.37 (adjusted *p*-value = 6.3 × 10^−5^); the transcription of NM_001107718 encoding the 70 kDa variant 2 was repressed by PE with a log_2_FC = -0.45 (adjusted *p*-value = 1.6 × 10^−5^) and by SRF S103D expression with a log_2_FC = -0.32 (adjusted *p*-value = 4.0 × 10^−3^). We now show by qRT-PCR using primers that detect both variants that CDC14A mRNA levels were reduced by 29% at 24 hours of PE treatment (Figure 1B).

As CDC14A mRNA levels were reduced by αAR stimulation, we tested whether inhibition of CDC14A expression would phenocopy the effects of PE on myocyte morphology. CDC14A RNAi was conferred by transfection of myocytes with pooled CDC14A small interfering RNA (siCDC14A) oligonucleotides designed to inhibit all known and predicted rat CDC14A alternatively-spliced isoforms. The siCDC14A were validated by immunoprecipitation of CDC14A protein from control and CDC14A siRNA-transfected myocytes, in which a single ∼65 kDa protein band was detected only in siControl samples (Figure 1C). CDC14A siRNA increased the width of myocytes similarly to PE treatment, 9% and 10%, respectively (Figure 1D,E and Figure S1A; for reference, in mice there is ∼20% increase in cardiomyocyte width after 2 weeks of pressure overload^10,15^). Like PE, CDC14A siRNA transfection did not affect myocyte length, resulting in significantly increased W:L ratios. Conversely, adenovirus-mediated expression of recombinant CDC14A wild-type protein (Flag and ALFA-tagged human variant 1 homolog) prevented the increase in W:L ratio induced by PE, whereas having no effect in the absence of the αAR-agonist (Figure 1F and Figure S1B). In addition, a catalytically inactive CDC14A C278S mutant^16^ acted as a dominant negative, inducing agonist-independent increased myocyte width and W:L ratio like CDC14A siRNA. Consistent with the repression of *Cdc14A* transcription by SRF S103D expression, wild-type CDC14A over-expression also inhibited the increased myocyte width and W:L ratio due to SRF S103D expression, without affecting myocyte length (Figure 1G and Figure S1C). Together, these data suggest that CDC14A serves as a brake on the growth in width of cardiomyocytes *in vitro*, whereas decreased CDC14A expression promotes the increased myocyte W:L ratio induced by the αAR-ERK1/2-RSK3-SRF regulatory pathway.

### KMT5A – An Effector for CDC14A in Cardiomyocytes

A review of the literature identified the lysine methyltransferase KMT5A as a CDC14A substrate.^17^ Using pooled siRNA targeting all rat KMT5A isoforms, two KMT5A-specific protein bands of ∼41 and ∼33 kDa were identified in cardiomyocytes, corresponding to the molecular weight of known alternatively-spliced isoforms (Figure 2A,B). Consistent with previous findings that CDC14A-catalyzed KMT5A dephosphorylation induces KMT5A nuclear export, ubiquitination, and proteosome-mediated degradation,^17^ CDC14A siRNA elevated KMT5A protein levels in cardiomyocytes (Figure 2C), whereas CDC14A over-expression reduced KMT5A protein levels (Figure 2D). As decreased CDC14A increased KMT5A expression, we hypothesized that KMT5A would be required for the myocyte growth in width induced by αAR signaling. Depletion of KMT5A in myocytes by KMT5A siRNA had no significant effect on the morphology of unstimulated cells, whereas KMT5A siRNA inhibited the increase in width and W:L ratio induced by PE (Figure 2E and Figure S2A). Notably, KMT5A siRNA also reversed the hypertrophic effects of CDC14A siRNA (Figure 2F and Figure S2B).

**Figure 2.**
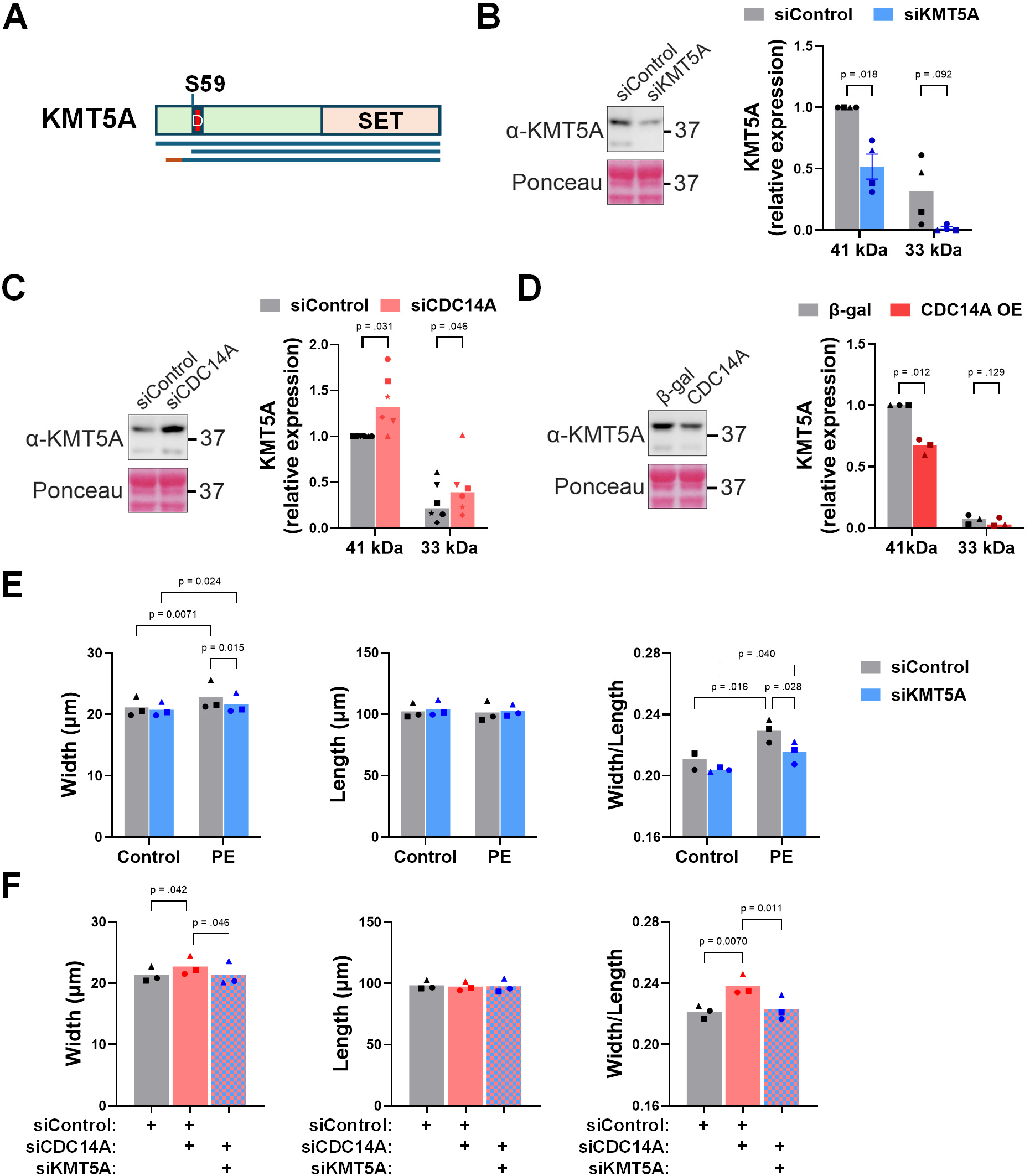
The CDC14A substrate KMT5A is required for the growth in width of cultured adult rat ventricular myocytes. **A.** KMT5A protein structure. The D-box (D) is an anaphase-promoting complex (APC) recognition motif required for ubiquitination and regulated by S59 phosphorylation.^17^ Blue and orange bars represent alternatively-spliced KMT5A isoforms. **B-D.** KMT5A (rabbit monoclonal antibody) Western blots detecting specific ∼41 and ∼33 kDa bands and Ponceau total protein stain of adult rat ventricular myocyte extracts following transfection with siRNA or adenoviral transduction to express Flag-CDC14A WT or β-galactosidase (β-gal) control. Representative blots shown for *n =* 4 (B), 6 (C), 3 (D). Comparisons by paired t-tests. **E,F.** Myocytes transfected 24 hours with KMT5A, CDC14A, or control siRNA were treated as indicated for 24 hours ± 20 µmol/L phenylephrine (PE) before measurement of individual myocyte width and length (F – no agonist condition only). Mean values for 3 individual myocyte preparations are indicated using different symbols. Data (matched for myocyte preparation) analyzed by 2-way ANOVA and Uncorrected Fisher’s LSD test (E) and 1-way ANOVA and Tukey’s Post-hoc test (F). See Figure S2 for myocyte images.

CDC14A dephosphorylates KMT5A serine residue 59 (Figure 2A).^17^ As shown previously in HeLa cells,^17^ a S59D phosphomimetic myc-tagged KMT5A mutant protein was enriched in the nucleus of adult cardiomyocytes when compared to myc-tagged KMT5A wild-type and S59A mutant (Figure 3A). Remarkably, myc-tagged KMT5A S59D, but not KMT5A S59A increased myocyte W:L ratio, phenocopying PE treatment (Figure 3B and Figure S3A,B). These data suggest that nuclear KMT5A, repressed by CDC14A in unstimulated cells, is required for the selective growth in width of cardiomyocytes induced by αAR stimulation.

**Figure 3.**
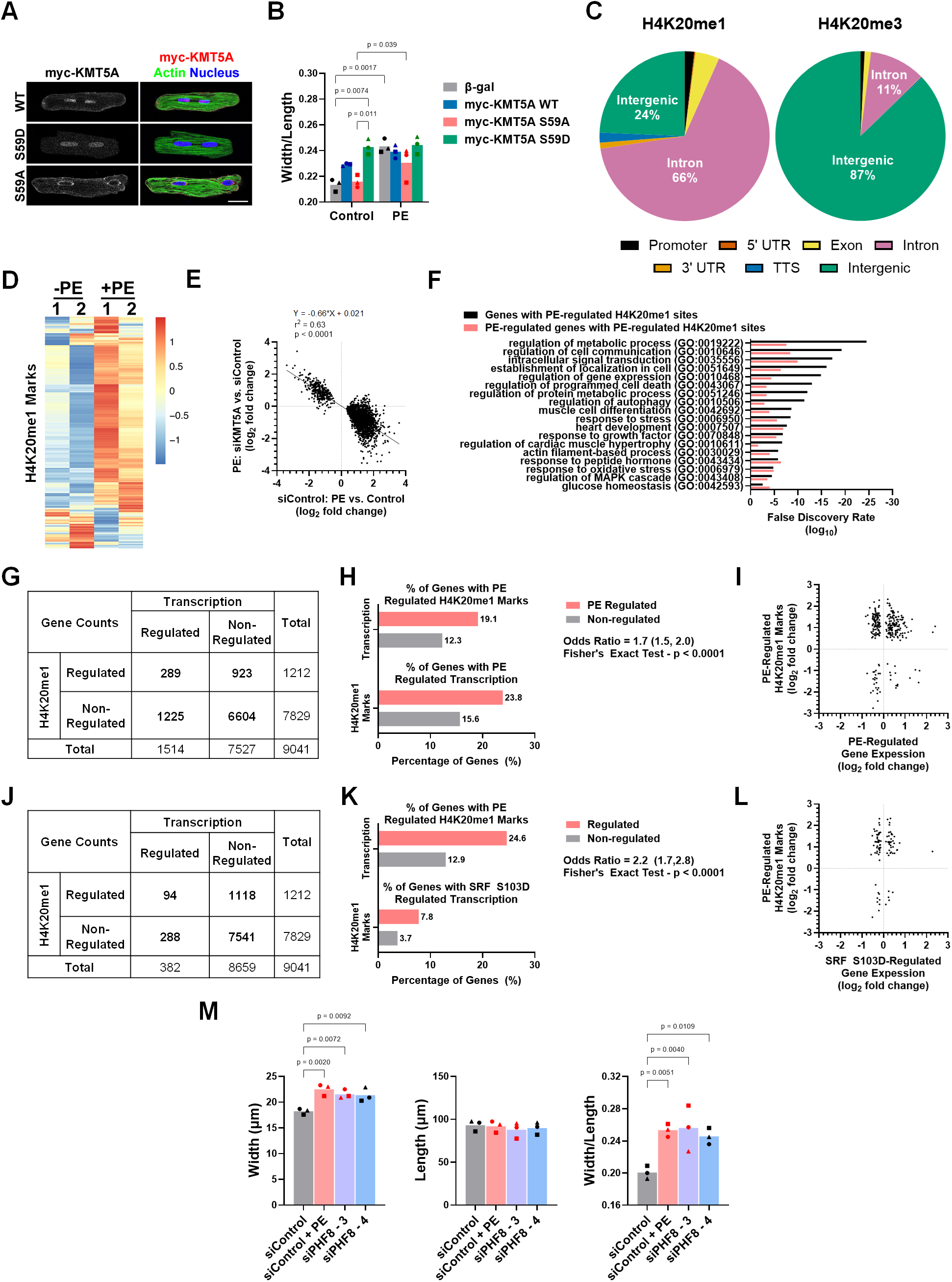
H4K20me1 and gene expression in adult rat myocyte hypertrophy. **A,B.** Nuclear KMT5A S59D increases myocyte W:L ratio. A - Myocytes were transduced with adenovirus for β-gal or myc-tagged KMT5A proteins for 2 days, then stained with myc-tag antibody (grayscale and red in composite images), phalloidin (Actin, green) and Dapi (Nucleus, blue). Scale bar – 20 µm. B - Additional myocytes were treated ± 20 µmol/L phenylephrine (PE) for 24 hours before measurement of individual myocyte width and length. Mean values for 3 individual myocyte preparations are indicated using different symbols. Data were analyzed by 2-way ANOVA (matched for myocyte preparation) and Tukey’s Post-hoc test. See Figure S3A,B for representative images and myocyte width and length data. **C.** CUT&RUN whole genome analysis was performed using H4K20me1 and H4K20me3 antibodies (n = 2 independent preparations). The distribution of marked sites on the rat genome was determined, distinguishing sites on promoters, 5’ untranslated regions (5’ UTR), coding exons, introns, 3’ untranslated regions (3’ UTR), transcriptional termination sites (TTS), and intergenic regions. In this study, “intragenic” sites include those on transcribed sequences, i.e. 5’UTRs, exons, introns, and 3’ UTRs. **D.** Heatmap of 2368 H4K20me1 sites regulated by PE (20 µmol/L for 1 h). **E.** Linear regression of the effect of KMT5A siRNA on PE-treated adult myocytes versus the effect of PE on control siRNA-transfected myocytes for the H4K20me1 marks shown in D. **F.** Gene ontology analysis for 1245 Genes containing (intragenic) PE-regulated H4K20me1 sites. Complete ontology list shown in Table S3. **G-H.** Contingency analysis of the 9041 transcripts containing Intragenic H4K20me1 sites comparing PE-regulation of gene transcription (Precision Nuclear Run-On Sequencing [PRO-seq], NCBI GEO accession number GSE134801) versus PE-regulation of H4K20me marks. **I.** Comparison of PE-induced changes in gene transcription and H4K20me1 marks for the genes and sites in G-H. **J-L.** Same as G-I for SRF S103D-regulated gene transcription. **M.** Myocytes transfected 24 hours with siPHF8 siRNA (sequence 3 or 4) or control siRNA were treated as indicated for 24 hours ± 20 µmol/L PE before measurement of individual myocyte width and length. Mean values for 3 individual myocyte preparations are indicated using different symbols. Data were analyzed by 1-way ANOVA (matched for myocyte preparation) with Dunnett’s multiple comparison test for differences from siControl. See Figure S3C for representative images.

### Epigenomic Regulation by H4K20me1 and Hypertrophic Signaling

Of the known KMT5A substrates, histone 4 lysine residue 20 (H4K20) is the best characterized and represents a candidate mechanism by which nuclear KMT5A regulates cardiac myocyte morphology.^18^ KMT5A is the only known monomethyl transferase for H4K20, whereas subsequent di- and tri-methylation of H4K20 is catalyzed by other methyltransferases.^19^ As previously shown in other cell types,^20^ we found by CUT&RUN whole genome site mapping that in cardiomyocytes H4K20me1 marks were enriched (72%) in transcribed intragenic chromatin (5’ and 3’ untranslated regions, other exonic sequence, and introns), especially introns (66%), whereas H4K20me3 marked sites were enriched (87%) in intergenic chromatin (Figure 3C and Table S1). 2368 H4K20me1 marked sites (2.3% of the total 101844 sites detected by CUT&RUN) were regulated by PE treatment in adult myocytes (*p* < 0.05, Figure 4D). 87% of these regulated marks were increased by PE treatment. Comparison of the difference in H4K20me1 marks in PE-treated cells transfected with KMT5A or control siRNA (Figure 3E, y-axis) versus the effect on H4K20me1 markings by PE-treatment (x-axis) confirmed that for most sites the PE-regulation of H4K20me1 marking was KMT5A dependent.

**Figure 4.**
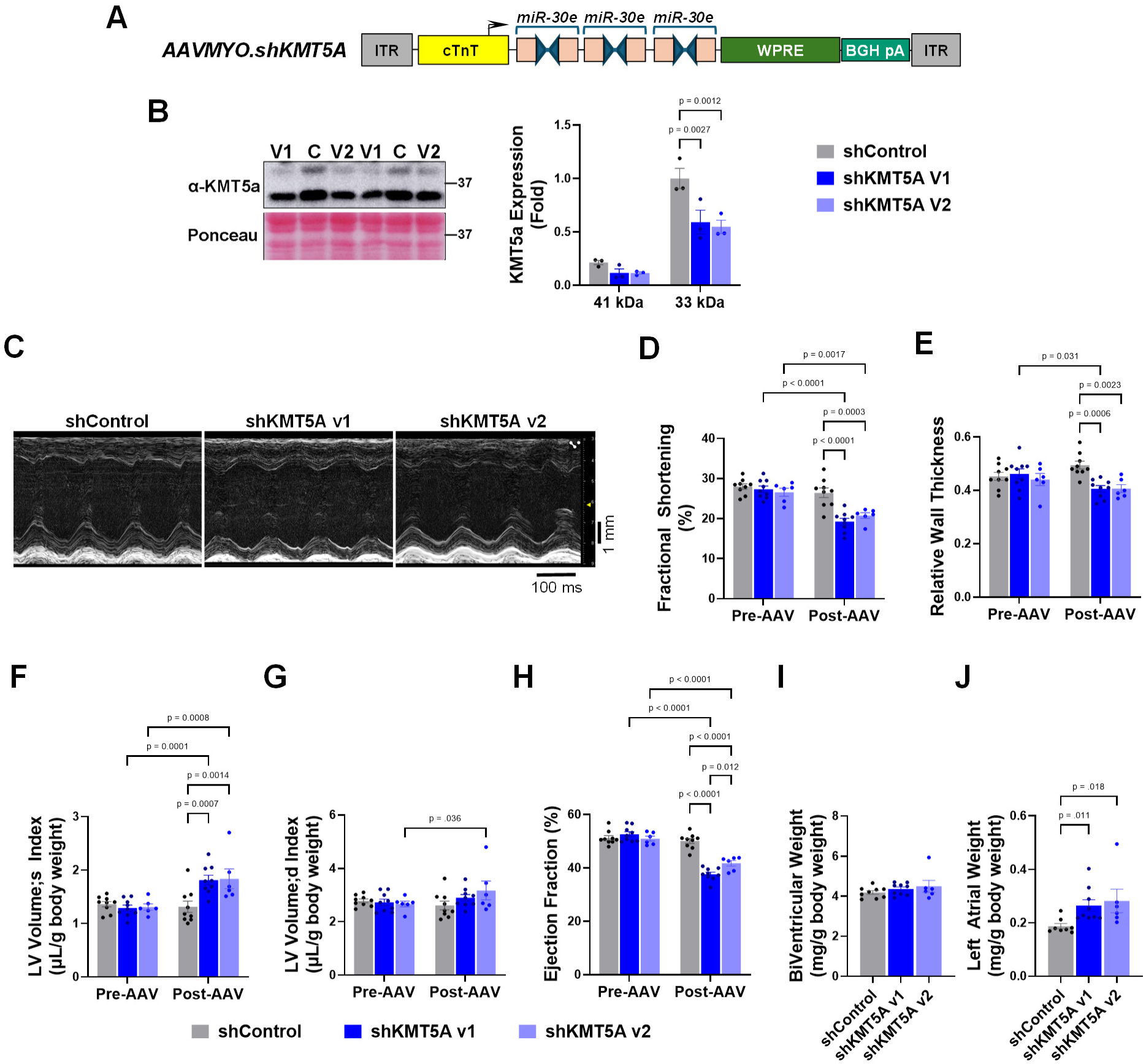
Inhibited KMT5A expression in mice induces cardiac systolic dysfunction and eccentric remodeling. **A.** Genome structure for adeno-associated virus (AAV) vectors containing the myotropic AAVMYO capsid^13^ and a cardiomyocyte-specific chicken troponin T promoter (cTnT)^28^. Two versions (V1 and V2) were constructed, each containing 3 mir-30e shRNA cassettes targeting 6 different mouse KMT5A sequences. ITR – inverted terminal repeat; WPRE - Woodchuck Hepatitis Virus Posttranscriptional Regulatory Element; BGH pA – bovine growth hormone polyadenylation sequence. **B.** KMT5A (rabbit monoclonal antibody) Western blots detecting specific ∼41 and ∼33 kDa bands and Ponceau total protein stain of left ventricular (LV) extracts from mice 3 months after intravenous administration of 10^11^ viral genomes AAVMYO.shKMT5A V1 or V2 or AAVMYO.shControl. Data analyzed by two-way ANOVA (matched by mouse) and Tukey’s multiple comparison test. *n* = 3 per cohort. **C-I.** 6-week-old male C57BL/6NJ mice were randomized by echocardiography (Pre-AAV) and administered AAV as in B with endpoint studies 1 month after AAV infusion (Post-AAV). **C.** Representative M-mode echocardiography at endpoint. **D, E.** Fractional shortening and relative wall thickness by M-mode echocardiography. Relative Wall Thickness = (PW + AW)/ID in diastole where PW, AW, and ID are LV posterior and anterior wall thickness and interior diameter. **F-H.** LV volumes and ejection fraction by 4-dimensional echocardiography. See Videos S1-S3. **I,J.** Gravimetric analysis of biventricular and left atrial weight indexed to body weight at endpoint. D-H - Data analyzed by 2-way ANOVA (matched for pre- and post-treatment) and Tukey’s multiple comparison test. I,J - Data analyzed by Kruskal-Wallis test with Dunn’s multiple comparison test. *n:* AAVMYO.shKMT5A V1 – 9; AAVMYO.shKMT5A V2 – 6; AAVMYO.shControl – 9 (D-I), 8 (J).

To study how H4K20me1 marks affect gene expression, marked sites were mapped to the rat reference sequence genome rn7. The intragenic H4K20me1 sites mapped to 9249 unique transcripts (Table S1, 99% of marks successfully mapped), of which 9041 (98%) were quantitated in the earlier PRO-seq study of *de novo* transcription (Table S2).^11^ 1245 transcripts (13% of the total 9249) contained at least one PE-regulated H4K20me1 mark. Gene ontology analysis showed that this subset of transcripts with PE-regulated H4K20me1 marks were enriched in genes regulating metabolism, intracellular signaling, cell death, and cardiac development and hypertrophy (Figure 3F and Table S3). A limited survey of the literature pertaining to the genes with regulated marks identified many known to be involved in cardiac remodeling or heart failure (Table S1), including, for example, *Akap6*, *Atp2a2 (*sarcoplasmic/endoplasmic reticulum calcium ATPase 2*), Mapk1 (*extracellular signal-regulated kinase *2), Nppa* (atrial natriuretic factor), and *Ppp3cb (*calcineurin Aβ).^1^ Notably, genes containing PE-regulated H4K20me1 marks included *Bag3*, *Cacna1c, Flnc*, *Jph2*, *Klhl24*, *Ldb3*, *Mybpc3*, *Plekhm2*, *Rbm20,* and *Vcl,* whose mutation in patients is associated with Dilated or Hypertrophic Cardiomyopathy.^21,22^

Comparison of the CUT&RUN and PRO-seq datasets showed that 289 transcripts were regulated by PE both in transcription and in H4K20me1 markings (Table S2). Similar gene ontologies were associated with this smaller group of transcripts (Figure 3F and Table S4). Remarkably, contingency analysis showed that PE-regulated H4K20m1 marking and gene transcription were associated with an Odds Ratio = 1.7, demonstrating a significant association between altered epigenomic marking and gene expression (Figure 3G,H). Increased H4K20me1 marks have been associated with both gene activation and repression, apparently promoting an open chromatin conformation permissive for dynamic gene regulation.^23,24^ Accordingly, we found altered H4K20me1 marking was not predictive of PE-induced gene repression or activation (Figure 3I). We hypothesize that via CDC14A reduced expression, KMT5A is increased in expression downstream of αAR and SRF activation. Like PE-regulated transcription, SRF S103D-induced transcription was associated with PE-regulated H4K20me1 markings (Odds Ratio = 2.2, Figure 3J-L). Notably, 55 genes were significantly regulated by both PE and SRF S103D in the PRO-seq analysis and in the H4K20me1 CUT&RUN analysis. These genes included *Rbms1* and *Mef2a,* which can promote pathological cardiac hypertrophy and systolic dysfunction and were transcriptionally downregulated by PE and SRF S103D,^25,26^ as well as the aforementioned cardiomyopathy-associated genes *Cacna1c*, *Flnc, Ldb3*, and *Mybpc3* (Table S2).^21,22^ Taken together, these results support the hypothesis that KMT5A-dependent H4K20me1 epigenomic modification contributes to the hypertrophic gene expression program induced by αAR and SRF activation.

To obtain independent evidence for a function of H4K20me1 in regulating myocyte growth in width, we used a second approach not relying on KMT5A gain or loss of function. Whereas KMT5A catalyzes H4K20 mono-methylation, Plant Homeodomain Finger Protein 8 (PHF8, KDM57B) is a demethylase for H4K20me1.^27^ Inhibited PHF8 expression by transfection with PHF8 siRNA increased adult myocyte width and W:L ratio comparably to αAR stimulation (Figures 3M and S3C). In sum, these results suggest that increased H4K20 mono-methylation promotes gene expression driving myocyte preferential growth in width.

### Inhibited KMT5A Expression Induces Eccentric Cardiac Remodeling

Both hypertrophy assays and epigenomic studies *in vitro* suggested that KMT5A plays an important role in regulating myocyte hypertrophy and morphology. To test directly the relevance of KMT5A to cardiac physiology, we expressed KMT5A shRNA *in vivo* using two different AAV vectors (V1 and V2) selective for the cardiomyocyte by inclusion of AAVMYO capsid and the chicken cardiac troponin T promoter,^13,28^ each vector containing three different shKMT5A sequences (Figure 4A). Western blot of whole mouse heart extracts using KMT5A antibody detected bands of similar sizes to those observed in cultured adult rat cardiomyocytes (Figures 2B and 4B), albeit the relative ratio of the ∼41 and ∼33 kDa bands was opposite between the two sets of samples. Intravenous administration of AAVMYO.shKMT5A V1 and V2 resulted in 41-46% decreased KMT5A expression in whole heart extracts when compared to mice injected with AAVMYO.shControl virus (Figure 4B).

C57BL/6NJ mice were randomized for intravenous administration of AAVMYO.shKMT5A V1 or V2 viruses or AAVMYO.shControl virus and studied 1 month later. Results were similar for the two AAVMYO.shKMT5A viruses. M-mode echocardiography showed that inhibited KMT5A expression resulted in significant systolic dysfunction, including decreased fractional shortening (Figure 4D) and increased end-systolic LV interior diameter, with wall thinning and decreased relative wall thickness (Figure 4E and Figure S4). 4-dimensional imaging confirmed the shKMT5A-induced systolic dysfunction (Videos S1-S3). Indexed end-systolic LV volume was increased by 38-40%, and LV ejection fraction was 8-12% lower following AAVMYO.shKMT5A V1 and V2 injection, with minimal evidence of LV dilatation (Figure 4E-G). One month after AAV administration, there was no significant ventricular hypertrophy (Figure 4H). However, KMT5A RNAi did result in left atrial hypertrophy (41-50% increased indexed left atrial weight, Figure 4I), reflective of either a particularly important role for KMT5A in atrial physiology or atrial hypertrophy secondary to LV dysfunction. Taken together, these results show that KMT5A is important for the maintenance of physiological cardiac structure and function. Moreover, the induction by shKMT5A of eccentric remodeling was consistent with observations *in vitro* that KMT5A promotes increased cardiomyocyte W:L ratio.

### Targeting of CDC14A-KMT5A as a Treatment for Dilated Cardiomyopathy

As DCM is associated with eccentric cardiac hypertrophy and decreased myocyte W:L ratio,^4,29,30^ we hypothesized that KMT5A would be reduced in DCM (at least for some etiologies), such that inhibiting CDC14A would improve cardiac structure and function in DCM via restoration of KMT5A expression. The TM54 FVB/N mouse contains a mutant α-tropomyosin E54K transgene expressed under the control of the cardiac myocyte-specific α-myosin heavy chain promoter and develops a DCM phenotype within a month of age.^14,29^ Western blot analysis of whole heart extracts from 4-month-old TM54 and non-transgenic (NTG) wild-type littermates showed that the major ∼33 kDa KMT5A band was reduced 24% in TM54 mice (Figure 5A). CDC14A RNAi was induced *in vivo* using the gene therapy vector AAV9sc.shCDC14A that directs cardiomyocyte-selective expression of two CDC14A shRNAs targeting the N-terminal CDC14A catalytic domain (Figure 5B). Like *in vitro*, inhibited CDC14A expression increased the expression of both KMT5A bands in whole heart extracts 41-42% (Figure 5C-E).

**Figure 5.**
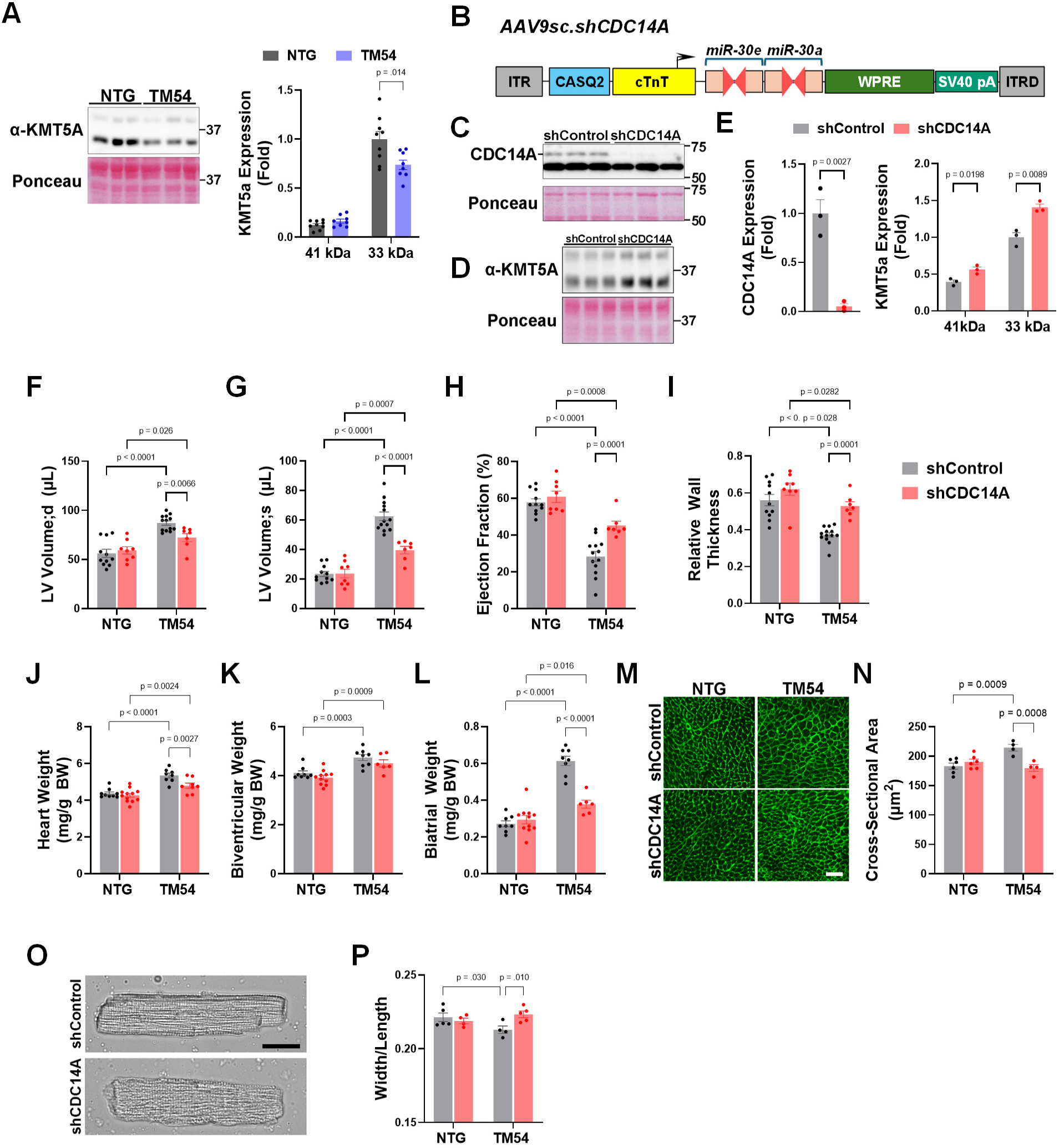
Inhibited CDC14A expression improves cardiac structure and function in a mouse model of Dilated Cardiomyopathy. **A.** Western blot using rabbit KMT5A antibodies and total Ponceau stain of left ventricular extracts from 4-month-old TM54 and non-transgenic (NTG) mice. unpaired *t*-test; *n* = 9 (NTG), 8 (TM54). **B.** AAV9sc.shCDC14A gene therapy vector contains two miR-30 mouse CDC14A shRNA cassettes expressed under the control of a calsequestrin (CASQ2) enhancer and a chicken cardiac troponin T (cTnT) promoter.^28,54^ **C-N.** 2- to 3-day-old TM54 and NTG mice of both sexes were injected IP with 10^11^ vg AAV9sc.shCDC14A or AAV9sc.shControl. **C-E.** Western blot using HRP-conjugated mouse anti-CDC14A and rabbit KMT5A antibodies and total Ponceau stains of LV extracts from 12-week-old TM54 mice. *n* = 3 per cohort. Data analyzed by t-tests. **F-H.** Echocardiography of 16-week-old mice for LV volumes and ejection fraction by B-mode. **I.** Relative Wall Thickness of 16-week-old mice = (PW + AW)/ID in diastole where PW, AW, and ID are LV posterior and anterior wall thickness and interior diameter by M-mode. *n* (F-I): NTG shControl – 11; TM54 shControl – 13; NTG shCDC14A – 8; TM54 shCDC14A – 7. **J-L.** Gravimetric analysis of 12-week-old mouse hearts indexed to body weight. *n*: NTG shControl – 9 (J), 8 (K,L); TM54 shControl – 8; NTG shCDC14A – 12 (J), 11 (K,L); TM54 shCDC14A – 8 (J), 6 (K,L). **M,N.** Wheat germ-agglutinin-stained LV tissue and quantification of myocyte cross-section area. Bar – 50 µm. *n*: NTG - 6; TM54 - 4. **O,P.** Morphometry of isolated ventricular myocytes showing improved width:length ratio. Representative TM54 images shown. Bar – 25 µm. *n*: NTG shControl – 5; TM54 shControl – 4; NTG shCDC14A – 4; TM54 shCDC14A – 5. Data in F-P analyzed by 2-way ANOVA with post-hoc testing by Uncorrected Fisher’s LSD.

AAV9sc.shCDC14A or control AAV9sc.shControl was administered to neonatal TM54 and NTG mice. CDC14A RNAi had no significant effect on wild-type NTG mice. At 16 weeks of age, AAV9sc.shCDC14A-treated TM54 mice exhibited improved left ventricular (LV) volumes (both diastole and systole, Figure 5F,G) and a 17% increase in ejection fraction by B-mode echocardiography (shControl - 28.3±2.8% vs. shCDC14A – 45.1±2.4%, *p* < 0.0001; Figure 5H and Figure S5A,B). M-mode echocardiography corroborated the improvement in systolic function (fractional shortening) and demonstrated increased LV wall thicknesses and relative wall thickness for AAV9sc.shCDC14A-treated TM54 mice (Figure 5I and Figure S5C,D). Cardiac hypertrophy was attenuated by AAV9sc.shCDC14A, with atrial hypertrophy inhibited 77% by CDC14A RNAi (Figure 5J-L).

The decrease in LV absolute and relative wall thickness in DCM results from side-to-side slippage and loss of myocytes and decreased myocyte W:L ratio.^30^ However, individual DCM myocytes typically exhibit hypertrophy with both increased width and length (i.e., with an increase in width less than in length).^29^ Wheat germ-agglutinin staining of tissue sections showed that LV cardiomyocyte cross-section area was increased in AAV9sc.shControl but not AAV9sc.shCDC14A transduced TM54 mice (Figure 5M,N). Importantly, morphometric analysis of isolated cardiomyocytes showed that, as expected,^29^ TM54 DCM was associated with decreased cardiomyocyte W:L ratio (Figure 5O,P and Figure S5E). Inhibition of CDC14A expression in TM54 mice normalized the myocyte W:L ratio. Neither the TM54 transgene nor CDC14A expression was associated with an altered number of nuclei per myocyte, including no increase in mononuclear cells as might be associated with increased cellular proliferation (Figure S5F). Finally, consistent with the improvement in cardiac remodeling, AAV9sc.shCDC14A prevented the mild interstitial myocardial fibrosis associated with TM54 DCM (Figure S5G).

RNA-sequencing (RNA-seq) was performed on total RNA isolated from TM54 and NTG mouse LV tissue. 3830 genes were differentially expressed between TM54 and NTG mice injected with AAV9.shControl, while 4131 genes were differentially expressed between AAV9.shCdc14A and AAV9.shControl injected TM54 mice, of which 1840 genes were differentially expressed in both comparisons (adjusted p-value < 0.05, Table S5). Heatmap analysis of the 1840 genes showed a remarkable normalization of TM54-altered gene expression by the AAV9.shCDC14A gene therapy (Figure 6A). Accordingly, gene expression changes due to AAV9.shCDC14A administration to TM54 mice were inversely correlated to those due to TM54 transgene expression (Figure 6B, r = [-0.85,-0.83]). Expanding the comparison to include all 3830 genes differentially expressed in AAV9.shControl TM54 and NTG mice, gene expression changes due to AAV9.shCDC14A administration to TM54 mice were still robustly inversely correlated to that due to TM54 transgene expression (r = [-0.69,-0.65]). Gene ontology analysis of the 1840 differentially expressed genes for both transgene and shCDC14A expression showed enrichment for genes related to metabolic pathways, intracellular signaling, and heart development (Figure 6C, Table S6). For example, *Srf* expression was decreased in TM54 mice and restored by shCDC14A, consistent with *Srf* serving as a marker of eccentric vs. concentric remodeling^11^ and suggesting potential feedback regulation of *Srf* transcription by CDC14A. In addition, like the results of H4K20me1 CUT&RUN analysis, several genes whose mutation in patients is associated with cardiomyopathy were normalized in expression by AAV9.shCDC14A, including *Flnc*, *Myl2*, *Jph2*, *Nkx2-5*, *Plekhm2*, *Pln*, and *Scn5a*.^21,22^ Further, 160 of the 1840 genes regulated in the TM54 study were also found *in vitro* to have PE-regulated H4K20me1 sites by CUT&RUN. Like the larger set of 1840 genes, this subset showed a dramatic reversal of gene expression by shCDC14A expression (Figure 6D), moreover, resulting in a stronger inverse correlation between gene expression changes due to AAV9.shCDC14A treatment and those due to TM54 disease induction (Figure 6E, r = [-0.94,-0.90]). Thus, analysis of gene expression corroborated at the molecular level the improvement in cardiac structure and function conferred by shCDC14A expression in the TM54 DCM mouse.

**Figure 6.**
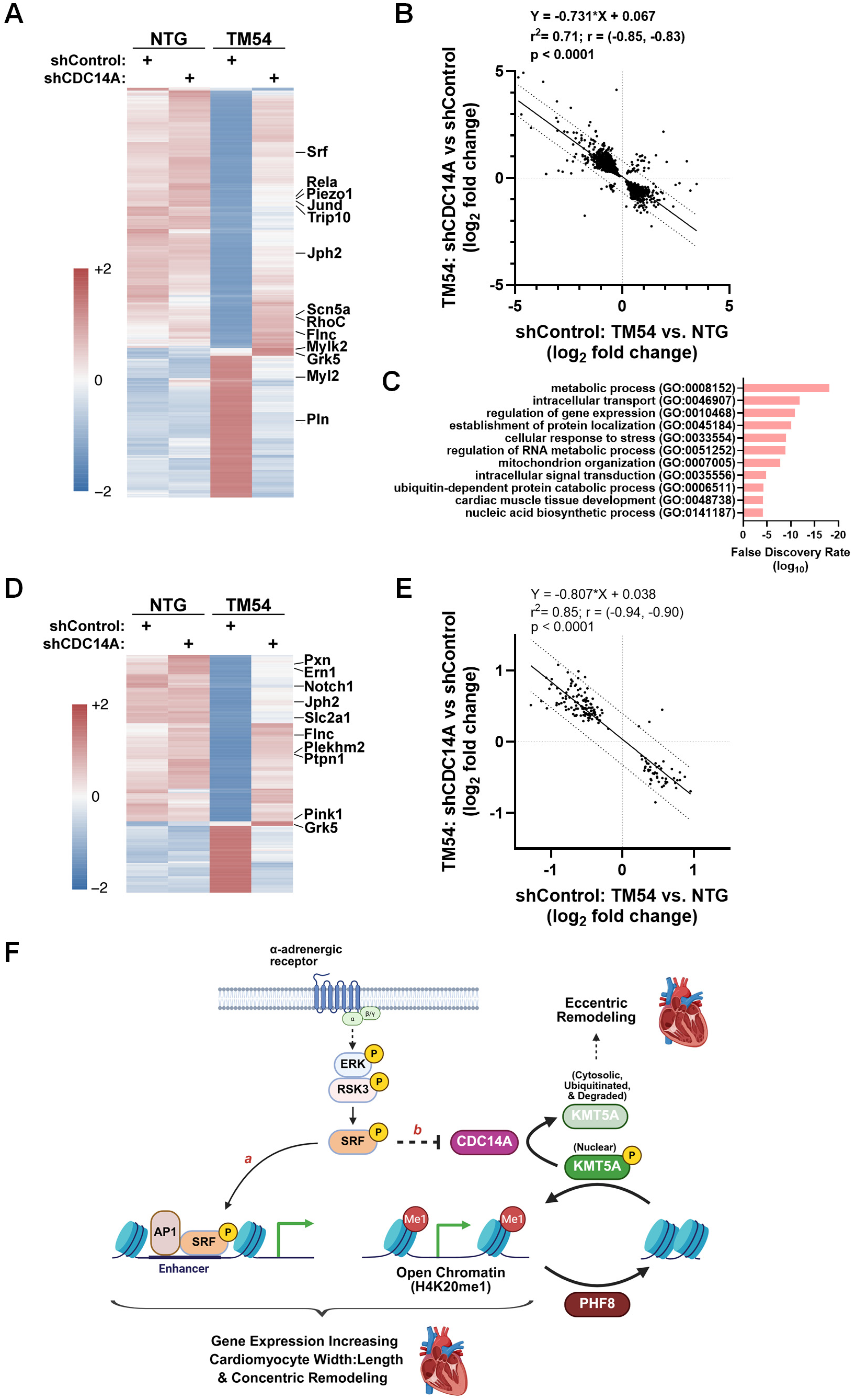
Inhibited CDC14A expression normalized cardiac gene expression in a mouse model of Dilated Cardiomyopathy. **A.** Heatmap of the 1840 genes differentially expressed in the left ventricle both between TM54 and non-transgenic (NTG) mice injected with AAV9.shControl and between AAV9.shCdc14A and AAV9.shControl injected TM54 mice (adjusted p-value < 0.05, Table S5). Average values for cohorts are shown. *n* = NTG shControl – 3; TM54 shControl – 6; NTG shCDC14A – 4; TM54 shCDC14A – 4. **B.** Linear regression of log2 fold change for the effect of shCDC14A on TM54 LV gene expression versus the effect of the TM54 transgene on LV gene expression in shControl mice. Pearson r 95% confidence interval and p-value provided. **C.** Gene ontologies for genes in A. Complete list in Table S6. **D,E.** Same as A,B for the 160 genes also identified as containing PE-regulated H4K20me1 in cultured adult myocytes by CUT&RUN (see Figure 3). **F.** Model for regulation of cardiac remodeling by the ERK/RSK3/SRF pathway. Ser-103 phosphorylated-SRF promotes concentric remodeling via dual mechanisms: a) as shown previously, phosphorylated SRF activates enhancers binding phosphorylated SRF and AP1 transcription factors, inducing gene expression promoting remodeling characterized by myocyte preferential growth in width;^11^ b) as shown here, inhibited CDC14A expression permits Ser-59 phosphorylated-KMT5A to increase H4K20me1 epigenomic modification, inducing an open chromatin conformation and facilitating gene expression promoting remodeling characterized by myocyte preferential growth in width. Opposing this process, CDC14A and the demethylase PHF8 inhibit H4K20 mono-methylation and promote eccentric cardiac remodeling. Created in BioRender. Kapiloff, M. (2026) https://BioRender.com/64vlxjk

## Discussion

In this study, we describe a new signaling pathway regulating myocyte morphology and the balance between concentric and eccentric remodeling. Evidence is provided by gain and loss of function *in vitro* and *in vivo* and by mechanistic studies of epigenomic regulation of gene expression. An initial proof-of-concept is provided for targeting of CDC14A-KMT5A signaling in the treatment of DCM.

We establish here a role for CDC14A in the stressed cardiomyocyte. Although CDC14 protein phosphatase was first identified in yeast for its role in mitosis, mammalian CDC14A has been shown to have other functions, including in cell adhesion and migration, microtubule and actin stress fiber regulation, DNA repair and transcriptional regulation.^12,16,31^ Accordingly, CDC14A has been associated with tubulin-containing structures like centrosomes, microtubules, and cilia and has been detected in the nucleus and at actin filaments of the leading edges of cultured cells.^16,32^ The relatively low level of CDC14A protein in cultured cardiomyocytes has thus far limited our ability to localize CDC14A within the myocyte (See Figure 1C), but its previous detection in the nuclei of other cell types is intriguing, particularly given CDC14A’s regulation of KMT5A and gene expression.^16^ *CDC14A* mutations have not been reported to be associated with cardiac defects, and shCDC14A expression had no apparent effect on baseline cardiac phenotype in mice (Figure 5). Instead, recessive *CDC14A* human mutations are associated with deafness and male infertility, traits that have been recapitulated in p.C278S knock-in mice lacking CDC14A catalytic activity.^16^ In mice, constitutive *Cdc14a* gene deletion and p.C278S mutation also result in partial perinatal lethality by an unknown mechanism.^16^

The current study suggests that in stressed cardiomyocytes CDC14A restrains concentric hypertrophy and promotes eccentric cardiac growth. Interestingly, in a microarray analysis of paired LV samples from LV assist device (LVAD) non-ischemic cardiomyopathy patients who improved sufficiently for LVAD explantation, CDC14A mRNA levels were reduced 2-fold at explant when compared to tissue obtained at LVAD implantation.^33^ However, elevated CDC14A mRNA levels do not appear to be a consistent marker for heart failure with reduced ejection fraction (HFrEF), as a meta-analysis of clinical heart failure studies reported variable CDC14A levels in ischemic and non-ischemic dilated cardiomyopathy (https://saezlab.shinyapps.io/reheat/).^34^ Accordingly, we found that *Cdc14a* was not significantly different in expression between TM54 and NTG mice (adjusted p-value = 0.58). This contrasts with SRF. SRF mRNA levels were reduced in TM54 mice (log2FC = -0.94, adjusted p-value = 0.0062) and in the meta-analysis of human heart failure (mean t = -1.24; fisher p-value = 1.5 × 10^−7^),^34^ in accord with our previous demonstration of reduced phosphorylated SRF in human heart failure tissue.^11^ Thus, while inhibition of CDC14A expression was beneficial in DCM, increased CDC14A expression is unlikely to be a common primary driver of HFrEF, and, instead, may act as a modifier of disease phenotype. These observations are consistent with a model in which, in conjunction with other factors, SRF activation indirectly represses *Cdc14a* gene transcription, especially since SRF S103D is associated with gene activation.^11^ Future studies will be necessary to define the upstream signals besides αAR stimulation and SRF that regulate CDC14A expression, as well as the likely regulation in cardiomyocytes of CDC14A by post-translational modification.^35^

CDC14 phosphatases catalyze the dephosphorylation of phosphoserine residues within the consensus site pSer-Pro-X-Lys/Arg.^36^ The list of known substrates for CDC14A is relatively limited, mainly proteins involved in the regulation of the cell cycle and the actin cytoskeleton.^12,37–39^ In this study, we focused on KMT5A as a CDC14A substrate. Interestingly, while KMT5A mRNA levels were not reduced in TM54 hearts (adjusted p-value = 0.92) and in some human heart failure studies,^19^ KMT5A mRNA levels were overall reduced in the aforementioned meta-analysis of human heart failure (mean t = -1.72; fisher p-value = 1.1 × 10^−6^).^34^ More importantly, KMT5A protein levels were reduced in TM54 mice (Figure 5A) and in a recently published human heart failure study,^40^ suggesting that decreased KMT5A protein may be a common feature of HFrEF.

In cycling cells, CDC14A/B catalyzes the dephosphorylation of KMT5A S59 (same as S29 in the 322 aa human isoform PR-Set7).^17^ In HeLa cells, S59 phosphorylation was shown to drive KMT5A nuclear localization, whereas S59 dephosphorylation promoted KMT5A nuclear export, association with anaphase promoting complex (APC) E3 ubiquitin ligase, and subsequent proteosomal degradation.^17^ Consistent with this proposed regulatory mechanism, we found that CDC14A negatively regulated KMT5A expression *in vitro* and *in vivo*. In addition, the KMT5A S59D mutant, which was nuclear localized, increased the W:L ratio of unstimulated cultured adult myocytes like siCDC14A. Recently, KMT5A protein translation was shown to be dependent in neonatal cardiomyocytes upon N4-acetylcytidine KMT5A mRNA modification catalyzed by NAT10 (N-acetyltransferase 10).^40^ Notably, NAT10 cardiomyocyte-specific knock-out in the mouse resulted in a DCM-like phenotype. The regulation of KMT5A by CDC14A and NAT10 suggests that KMT5A may serve to integrate multiple upstream signals regulating pathological gene expression.

A well characterized substrate for KMT5A is H4K20,^19^ and given the hypertrophic activity of the nuclear KMT5A S59D mutant, we focused on epigenomic modification in this project. H4K20me1 is an epigenetic mark in its own right, as well as the essential precursor for di- and trimethylated H4K20 catalyzed by other methyltransferases.^24^ Present at both transcriptionally active and inactive genes, H4K20me1 promotes an open chromatin conformation accessible to factors responsible for the dynamic regulation of gene transcription.^23,41,42^ Therefore, upon activation of enhancers in response to phenylephrine,^11^ for example, these H4K20me1 pre-marked transcription units, which are in a more open configuration, will be the gene targets that are best activated by the signal-dependent enhancers (Figure 6F). Further, H4K20me1 reduces RNA polymerase pausing during transcription.^43^ In contrast, H4K20me3 is associated with gene repression and chromatin compaction.^24^ With the exception of a report that H4K20me1 marks near the *Myh7* promoter were regulated by thyroid hormone in LV tissue,^44^ H4K20me1 has not apparently been studied in the heart. Key observations here include the striking difference in chromatin distribution between H4K20me1 and K4K20me3 sites and the association of elevated levels of H4K20 mono-methylation with transcriptional regulation by αAR stimulation and SRF active mutant expression. Most regulated H4K20me1 sites were intragenic and increased in marking by αAR stimulation, consistent with our proposed model for CDC14A-KMT5A signaling (Figure 6F). The 13% of sites decreased in H4K20me1 marks presumably reflect either site-specific induction of H4K20 di- or tri-methyltransferases or H4K20me1 demethylases. Notably, increased H4K20 mono-methylation was associated with PE- and SRF S103D-regulation of cardiomyocyte transcription (Figure 3), including the regulation of genes important for pathological cardiac remodeling and which are mutant in human cardiomyopathy. In addition, genes with αAR-regulated H4K20me1 marks *in vitro* were also regulated *in vivo* by CDC14A (Figure 6).

PHF8 is a demethylase for H4K20me1, as well as H3K9me1/2 and K3K27me2.^27^ Cardiomyocyte-specific PHF8 transgenic expression in mice has been shown to inhibit the development of pathological cardiac remodeling and heart failure in response to chronic pressure overload, while conferring no apparent phenotype in unstressed mice.^45^ In addition, PHF8 overexpression inhibited, whereas PHF8 RNAi potentiated PE-induced neonatal myocyte hypertrophy. Corroborating that KMT5A-dependent H4K20 mono-methylation regulates the genes that drive myocyte growth in width, we found that PHF8 RNAi also increased adult cardiomyocyte W:L ratio. Taken together, the results presented here define the CDC14A-KMT5A signaling pathway that regulates H4K20me1 epigenomic modification as a key co-regulator of cardiomyocyte gene expression that controls the balance between myocyte growth in length and width and between eccentric and concentric remodeling. We note, however, that other KMT5A substrates, namely DNMT1 (DNA methyltransferase 1),^46^ p53,^40,47^ SNIP1 (Smad Nuclear Interacting Protein 1),^48^ UHRF1 (Ubiquitin like with PHD and ring finger domains 1),^49^ and α-tubulin^50^ may also serve as effectors for CDC14A-KMT5A signaling in cardiomyocytes, which would be the subject for future studies.

Consistent with the inhibition of myocyte growth in width by KMT5A RNAi *in vitro*, shKMT5A expression *in vivo* resulted in adverse remodeling and systolic dysfunction. The interest in KMT5A-dependent methylation has largely arisen from studies of carcinogenesis, driving the development of KMT5A inhibitors as chemotherapeutic agents.^51^ However, the fact that shKMT5A expression rapidly induces a DCM-like phenotype in mice may complicate these efforts. In contrast, the improvement in cardiac structure and function in the TM54 mouse following shCDC14A gene therapy suggests that enhancing KMT5A activity may be beneficial in DCM. With a prevalence as high as 1 in 250 in the adult population and a relatively high mortality despite modern therapy, DCM comprises a significant cause of cardiovascular mortality and is the most common indication for heart transplantation.^52,53^ The development of effective therapies has been complicated by the diverse etiologies and large number of genes resulting in DCM.^21^ In this context, CDC14A and KMT5A may comprise etiology agnostic targets for DCM treatment, justifying further exploration of the translational potential of this newly identified cardiomyocyte regulatory pathway.

## Non-standard Abbreviations and Acronyms

AAV: adeno-associated virus
CDC14A: cell division cycle 14A
CUT&RUN: Cleavage Under Targets and Release Using Nuclease
DCM: Dilated Cardiomyopathy
ERK: extracellular signal-regulated kinase
GFP: green fluorescent protein
H4K20me1: mono-methylated histone 4 lysine 20
HFrEF: heart failure with reduced ejection fraction
KMT5A: lysine methyltransferase 5A
log_2_FC: log_2_ fold change
MAP-kinase: mitogen-activated protein kinase
PE: phenylephrine
PHF8: plant homeodomain finger protein 8
PRO-seq: Precision Nuclear Run-On Sequencing
RNAi: RNA interference
RNA-seq: RNA sequencing
SRF: serum response factor

## Acknowledgments

Drs. Xueyi, Li, Jinliang Li, Samuelsson, Nguyen, and Colombe: data collection, analysis and interpretation of experiments; Drs. Tan and Nair: bioinformatics analyses; Dr. Grimm: critical reagent; Drs. Kapiloff and Rosenfeld: project supervision, analysis and interpretation, writing and final approval of manuscript. This work utilized bioinformatics services and computing resources provided by the Stanford Genetics Bioinformatics Service Center (GBSC).

## Supplemental Material

Supplemental Methods

Major Resources Table

Figures S1-S5

Tables S1-S7

Videos S1-S3

References 55-59

